# Subgroup-Specific Associations of *GRIA* Genes Encoding AMPA Glutamate Receptor Subunits with Patient Survival in Medulloblastoma

**DOI:** 10.64898/2026.02.24.707769

**Authors:** Bruno Saciloto, Matheus Dalmolin, Julia Caroline Marcolin, Martina Lichtenfels, Julia Vanini, Isabella B. Schröder Roesler, Jurandir M. Ribas Filho, Osvaldo Malafaia, Marcelo A.C. Fernandes, Caroline Brunetto de Farias, Amanda Thomaz, Rafael Roesler, Gustavo R. Isolan

## Abstract

Brain cancers hijack biological systems involved in neural development and synaptic plasticity. Medulloblastoma (MB), the most common malignant brain tumor in children, is thought to arise from disruptions in neurodevelopmental programs. Glutamatergic transmission mediated by α-amino-3-hydroxy-5-methyl-4-isoxazolepropionic acid (AMPA) receptors (AMPARs) has been implicated in synaptic communication between adult brain tumors and surrounding neurons; however, the possible role of AMPARs in MB remains largely unexplored. Here, we analyzed the expression of genes encoding AMPAR subunits, *GRIA1–4*, in datasets of MB tumor and cell lines, revealing distinct expression patterns and associations with overall survival (OS) across molecular subgroups and histological variants. Expression levels differed among MB molecular subgroups. Analysis using single-cell RNA sequencing (scRNA-seq) was consistent with enrichment of *GRIA1* in Group 3 and *GRIA4* in SHH MB. Higher *GRIA1, GRIA2*, and *GRIA4* transcription was associated with more favorable patient outcomes in specific MB subgroups. In contrast, high expression of *GRIA3* in SHH, or of either *GRIA3* or *GRIA4* in Group 3 MB, was associated with worse prognosis. Particularly robust but opposing associations with patient survival were found for *GRIA3* and *GRIA4* in SHH MB. Analysis of *GRIA* mRNA levels in MB cell lines using both quantitative reverse transcription polymerase chain reaction (qRT-PCR) and data from The Human Protein Atlas, supported some of the gene expression patterns observed in tumors. Together, these findings suggest that *GRIA* genes and their corresponding AMPAR subunits may have subgroup-specific prognostic relevance in MB.

## Background

Fast excitatory neurotransmission in the central nervous system (CNS) is mediated predominantly by glutamate acting on α-amino-3-hydroxy-5-methyl-4-isoxazolepropionic acid (AMPA)–type ionotropic receptors (AMPARs). AMPARs are tetrameric ligand-gated ion channels assembled from four protein subunits, forming a pore that is primarily permeable to sodium ions. Binding of glutamate to AMPARs triggers channel opening, resulting in sodium influx and consequent depolarization of the postsynaptic neuronal membrane (Collingridge et al., 2009; Francis et al., 2025). The receptor subunits are encoded by four distinct genes located on different chromosomes: *GRIA1* (encoding GluA1, also known as GluR1), *GRIA2* (GluA2/GluR2), *GRIA3* (GluA3/GluR3), and *GRIA4* (GluA4/GluRA-D2) (Lu et al., 2009; Nakagawa, 2010; Salpietro et al., 2019; Wu et al., 2007). GluA2 is crucial in influencing AMPAR calcium permeability and channel conductance, whereas GluA1 drives long-term neural plasticity, and GluA4 is the rarest subunit in the adult brain but is particularly enriched in cerebellar interneurons (Dylan Hale et al., 2026; Emond et al., 2010; Henley & Wilkinson, 2016; Scrutton et al., 2025; Terashima et al., 2019).

A growing body of evidence implicates AMPAR-driven signaling in malignant brain tumors, especially glioblastoma (GBM). In this context, AMPARs have been shown to mediate functional synaptic connections between neurons and tumor cells, enabling GBM cells to integrate into neuronal circuits. These neuron–tumor synapses can promote tumor growth and highlight a critical role for glutamatergic signaling in neuro–tumoral interactions within the brain (Taylor et al., 2023; Venkataramani et al., 2019; Venkatesh et al., 2019).

Medulloblastoma (MB), the most common malignant brain tumor in children, is an embryonal neoplasm of the cerebellum that exemplifies cancer arising from disrupted neurodevelopmental programs. MB is currently classified into four principal molecular subgroups: Wingless (WNT)-activated, Sonic hedgehog (SHH)-activated, and the non-WNT/non-SHH categories, named Group 3 and Group 4 (Northcott et al., 2011; 2019; Taylor et al., 2012). These subgroups differ markedly in their developmental origins, oncogenic signaling pathways, and clinical behavior, and molecular classification has become central to risk stratification, therapeutic decision-making, and clinical trial design. WNT-activated MB is associated with a particularly favorable prognosis, whereas SHH-activated MB displays heterogeneous outcomes that are strongly influenced by the mutational status of *TP53*, with *TP53*-mutant tumors conferring high risk. Among all four subgroups, Group 3 MB is consistently associated with the poorest clinical outcomes (Juraschka & Taylor, 2019; Northcott et al, 2012a). In addition, extensive intra- and intertumoral heterogeneity has led to the identification of further molecular subtypes within these major groups (Cavalli et al., 2017; Northcott et al., 2012b; Schwalbe et al., 2017).

Consistent with its embryonal nature, MB arises from aberrations in cerebellar development affecting defined neural progenitor populations. WNT-activated MB originates from neuronal precursors located in the lower rhombic lip (RL), where oncogenic activation of WNT signaling occurs. This germinal zone contributes to the generation of mossy fiber and climbing fiber neurons within the brainstem nuclei. In contrast, SHH-activated MB derives from cerebellar granule cell precursors (GCPs), also referred to as granule neuron precursors or progenitors (GNPs), which arise from the upper RL and populate the external granular layer during development (Oliver et al., 2005; Wechsler-Reya & Scott, 1999; Yang et al., 2008). Group 3 and Group 4 MBs may share a developmental origin in glutamatergic progenitor populations of the fetal RL (Smith et al., 2022).

Taken together, these findings indicate that MB arises from progenitor cells with glutamatergic differentiation programs. *GRIA2* transcripts have been detected in the lower RL /external germinal layer during development. As RL-derived cerebellar granule neurons differentiate and form synapses, they express AMPA receptor subunits, prominently GluA4 during early development and GluA2 in mature neurons (Hagihara et al., 2011; Kita et al., 2021). Despite this developmental link, the contributions of glutamate receptors to MB biology and possible tumor–neuron interactions remain poorly understood. Messenger RNA (mRNA) expression of all AMPAR subunits has been demonstrated by quantitative reverse transcription polymerase chain reaction (qRT-PCR) in a small set of 6 MB samples, with *GRIA4* emerging as the most highly expressed subunit, followed by *GRIA3* (Brocke et al., 2010). In addition, both transcript and protein expression of GluA1 (*GRIA1*) have been reported in three MB-derived cell lines (Yoshioka et al., 1996).

Investigating changes in expression of genes encoding neurotransmitter and neurotrophin receptors within tumors may reveal novel candidate prognostic markers and therapeutic targets in MB (Fratini et al., 2023; Remke et al., 2013; Souza et al., 2022; Thomaz et al., 2020). Here, we report the expression patterns of *GRIA* genes and their association with patient survival across the distinct molecular subgroups of MB.

## Methods

### Study Population and Data Source

Gene expression data were obtained from the Gene Expression Omnibus (GEO) repository. The Pomaroy dataset established by Cho et al. (2011; accession number GSE202043) was used to compare MB tumors when all subgroups were combined (n = 194) with normal cerebellar tissue (n – 11). The dataset originally described by Cavalli et al. (2017; accession number GSE85217), comprising 763 primary MB samples classified into the four distinct molecular subgroups of MB, WNT, SHH, Group 3, and Group 4, was used for further comparisons. The tumors were also analyzed when divided into histological types as classic, desmoplastic/nodular (desmoplastic), large cell/anaplastic (LCA), and MB with extensive nodularity (MBEN).

### Microarray Data Processing and Gene Expression Analyses

Raw data derived from Affymetrix Human Gene 1.1 ST Arrays were acquired using the GEOquery R package. Data pre-processing involved background correction and log2 transformation to stabilize variance. Probe annotation was subsequently performed using the org.Hs.eg.db package to map probe IDs to HGNC gene symbols. For genes represented in multiple probes, expression values were aggregated by calculating the mean intensity. The analysis focused specifically on the expression levels of the four genes encoding AMPAR subunits, *GRIA1*, *GRIA2*, *GRIA3*, and *GRIA4*, in addition to a selected core set of genes encoding *N*-methyl-D-aspartate (NMDA) and kainate glutamate receptor subunits (NMDA, *GRIN1*, *GRIN2A*, *GRIN2B*; kainate, *GRIK1*, *GRIK2*, *GRIK3*, and *GRIK4)* in patient survival analyses. Comparisons of gene expression levels between MB tumors and cerebellar tissue were performed with the Mann-Whitney U test, and comparisons among MB molecular subgroups and histological subtypes were carried out using the Kruskal-Wallis test. The Benjamini-Hochberg (FDR) correction was used to adjust for multiple comparisons.

### Gene Expression in MB Single-Cell RNA Sequencing

Expression of *GRIA1*-*GRIA4* in the four MB subgroups at the single cell level was accessed using MB single-cell RNA sequencing (scRNA-seq) data available in the GEO database under the accession number GSE156053 (https://www.ncbi.nlm.nih.gov/geo/query/acc.cgi?acc=GSE156053; Riemondy et al., 2022). The complete dataset of MB scRNA-seq populations can be accessed through the interactive online resource provided by the Pediatric Neuro-oncology Cell Atlas (https://www.pneuroonccellatlas.org/). This allowed Uniform Manifold Approximation and Projection (UMAP) visualization of *GRIA1-GRIA4* expression across MB subgroups.

### Survival Analysis

Overall survival (OS) of MB patients was analyzed using clinical metadata available within the dataset. From the total number of 763 primary MB samples available in the dataset, 612 tumors were selected on the basis of having associated OS data. Optimal cutoff points for dichotomizing transcription into high and low gene expression groups were determined using maximally selected rank statistics via the surv_cutpoint function from the survminer package. Kaplan-Meier OS curves were estimated using the survival package, and differences in survival probabilities between groups were assessed using the log-rank test. Analyses were stratified by tumor molecular subgroup to evaluate prognostic associations. Benjamini-Hochberg (BH, FDR) correction was used to adjust for multiple comparisons. Associations were considered statistically significant when both raw and adjusted *P* values were < 0.05. All statistical analyses were performed within the R statistical environment (v4.x).

### Gene Expression in MB Cell Lines

Messenger RNA expression of *GRIA1-4* in DAOY (SHH) and D283 (Group 3/4) MB cell lines was analyzed by RT-qPCR. RNA was extracted from cells using the ReliaPrep™ RNA Miniprep System (Promega, Madison, WI, USA), in accordance with the manufacturer’s instructions and quantified in NanoDrop (Thermo Fisher Scientific). The cDNA was obtained using the High Capacity cDNA Reverse Transcription Kit (Applied Biosystems), also according to the manufacturer’s instructions. The mRNA expression levels of *GRIA1*, *GRIA2*, *GRIA3*, and *GRIA4* were quantified using PowerUp SYBR Green Master Mix (Applied Biosystems). Relative gene expression levels were normalized to actin beta (*ACTB*) and calculated using the 2⁻ΔCt method. Statistical analyses were performed using ΔCt values. The primer sequences used are shown in Table 1.

**Table 1.**
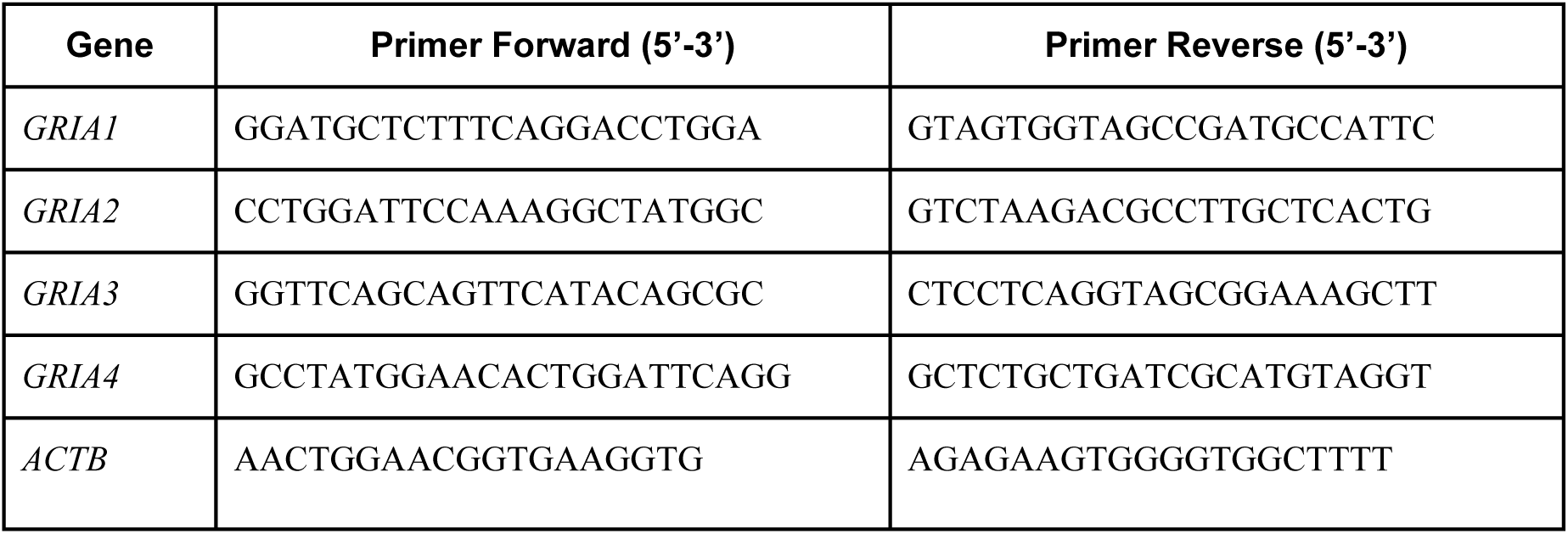
Primers used for RT-qPCR analysis of *GRIA1-4* expression.

Messenger RNA expression of *GRIA* genes in cell lines was also obtained from The Human Protein Atlas (https://www.proteinatlas.org/; DAOY (SHH) and D341 (Group 3) MB cell lines, *GRIA1*, https://www.proteinatlas.org/ENSG00000155511-GRIA1/cell+line#brain_cancer; *GRIA2*, https://www.proteinatlas.org/ENSG00000120251-GRIA2/cell+line#brain_cancer; *GRIA3*, https://www.proteinatlas.org/ENSG00000125675-GRIA3/cell+line#brain_cancer; *GRIA4*, https://www.proteinatlas.org/ENSG00000152578-GRIA4/cell+line#brain_cancer; NHAHTDD non-tumoral brain cells, *GRIA1*, https://www.proteinatlas.org/ENSG00000155511-GRIA1/cell+line#non-cancerous; *GRIA2*, https://www.proteinatlas.org/ENSG00000120251-GRIA2/cell+line#non-cancerous; *GRIA3*, https://www.proteinatlas.org/ENSG00000125675-GRIA3/cell+line#non-cancerous; *GRIA4*, https://www.proteinatlas.org/ENSG00000152578-GRIA4/cell+line#non-cancerous). Accessed on January 27^th^ 2026.

## Results

### Expression of *GRIA* Genes in MB Tumors and Normal Cerebellar Tissue

Levels of *GRIA1*, *GRIA3*, and *GRIA4*, but not *GRIA2*, were significantly lower in MB tumors in comparison with non-tumoral cerebellar tissue (Fig. 1).

**Fig 1.**
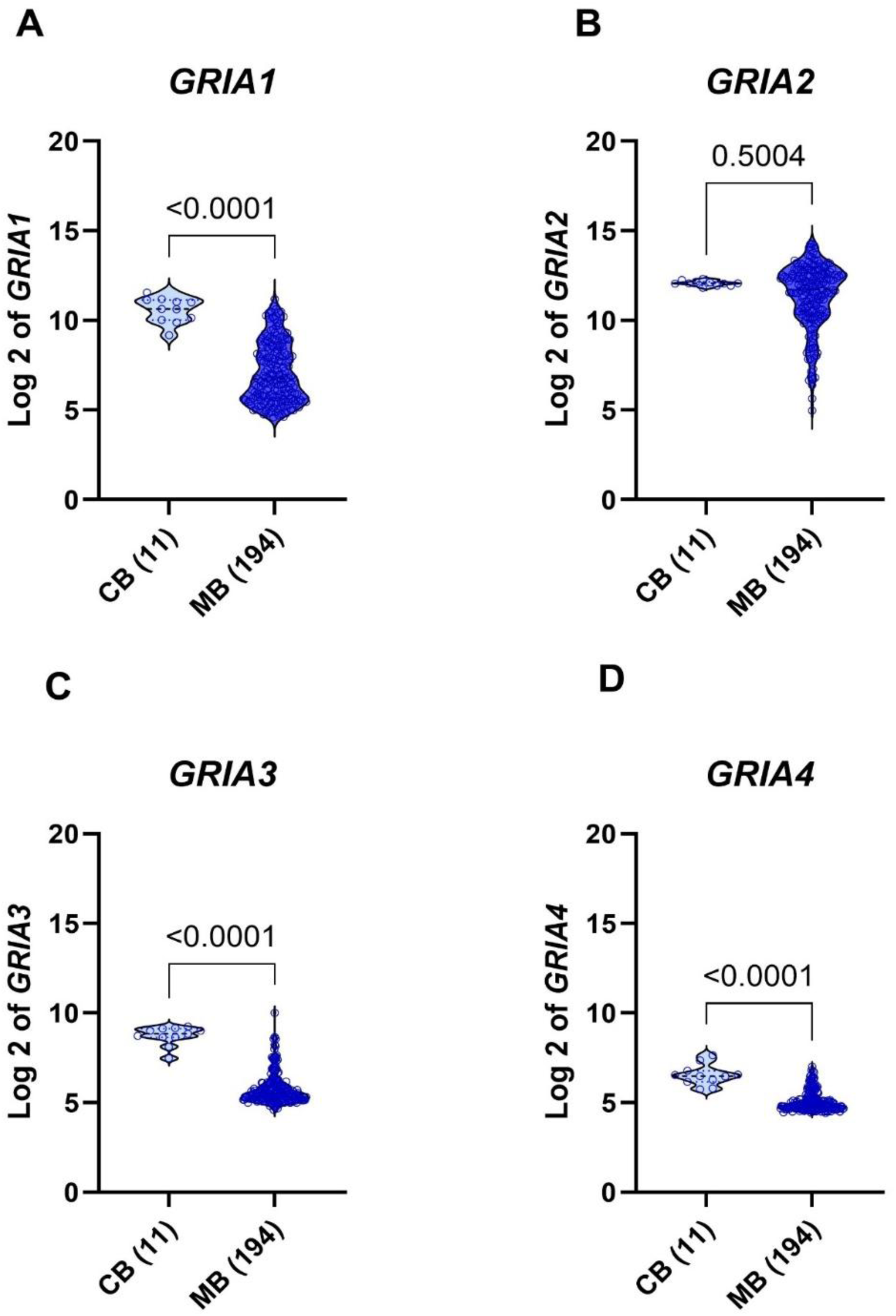
Gene expression levels of **A**, *GRIA1*, **B**, *GRIA2*, **C**, *GRIA3*, and **D**, *GRIA4* in MB tumors (n = 194) and normal cerebellar tissue (CB; n = 11). Data were obtained from the dataset established by Cho et al. (2011). Adjusted *P* values are indicated in the panels.

### Expression of *GRIA* Genes in MB Tumors Classified According to Molecular Subgroup

*GRIA1* transcription was higher in Group 3 MB compared to all other groups, whereas WNT and SHH tumors showed the lowest expression and did not differ from each other (Fig. 2A). Expression of *GRIA2* was higher in SHH and Group 4 MB and the lowest levels were observed in WNT MB (Fig. 2B). The only significant difference in *GRIA3* expression was found between WNT (lowest expression) and Group 4 (highest expression), although differences among groups were small for this gene (Fig. 2C). *GRIA4* expression was markedly higher in SHH compared to all other three subgroups (Fig. 2D).

**Fig 2.**
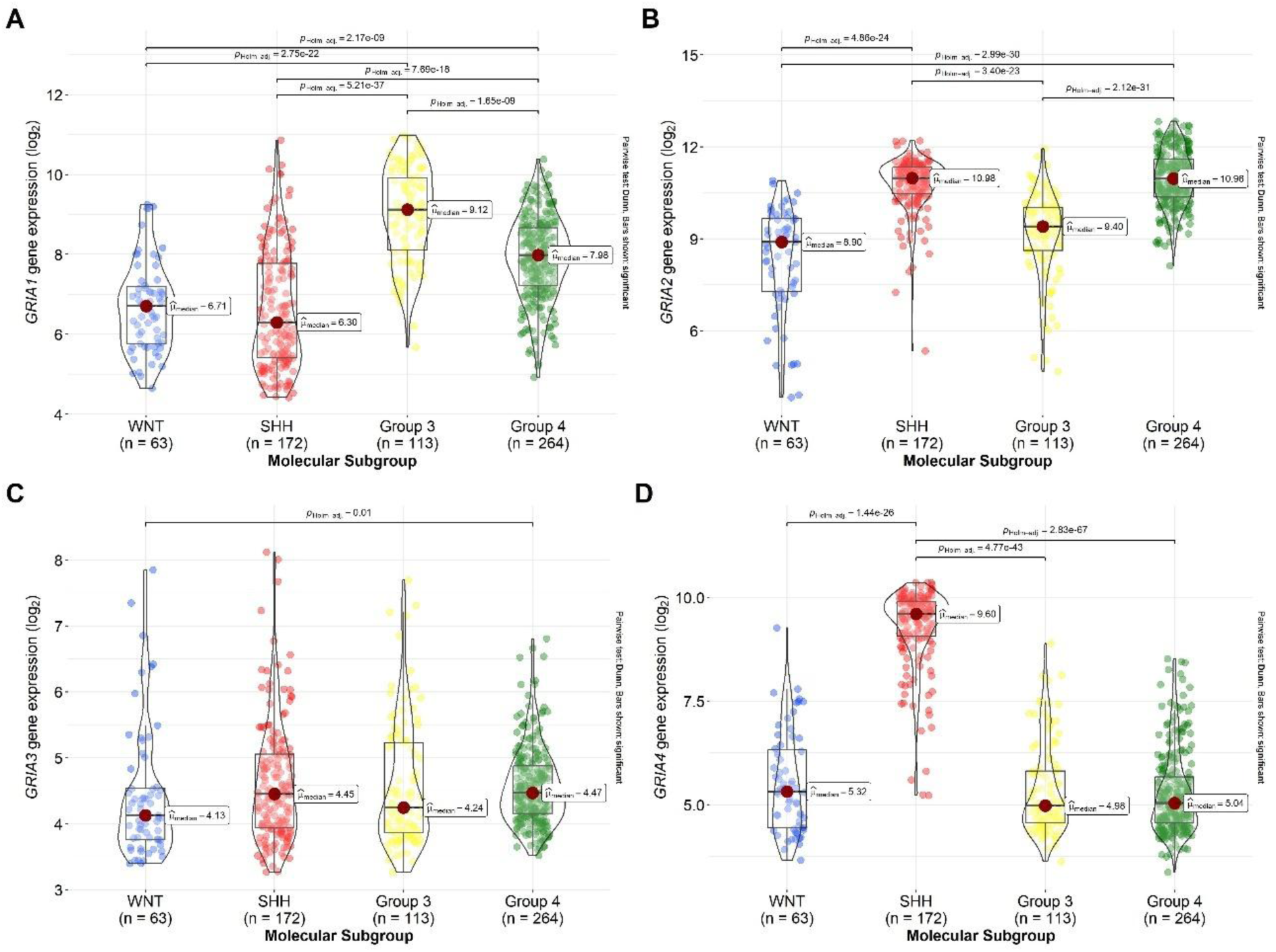
Gene expression levels of **A**, *GRIA1*, **B**, *GRIA2*, **C**, *GRIA3*, and **D**, *GRIA4* in MB tumors classified into the four molecular subgroups, WNT (n = 63), SHH (n = 172), Group 3 (n = 113), and Group 4 (n = 264). Data were obtained from the dataset established by Cavalli et al. (2017). Adjusted *P* values and significant differences are indicated in the panels.

### Expression of *GRIA* Genes in MB Tumors Classified According to Histological Type

Desmoplastic MB showed significantly lower levels of *GRIA1* compared to either classic or LCA tumors (Fig. 3A), and expression of both *GRIA2* and *GRIA4* was overall higher in desmoplastic and MBEN compared to classic and LCA MB (Fig. 3B, 3D). There was no significant difference among histological variants in *GRIA3* transcription (Fig. 3C).

**Fig 3.**
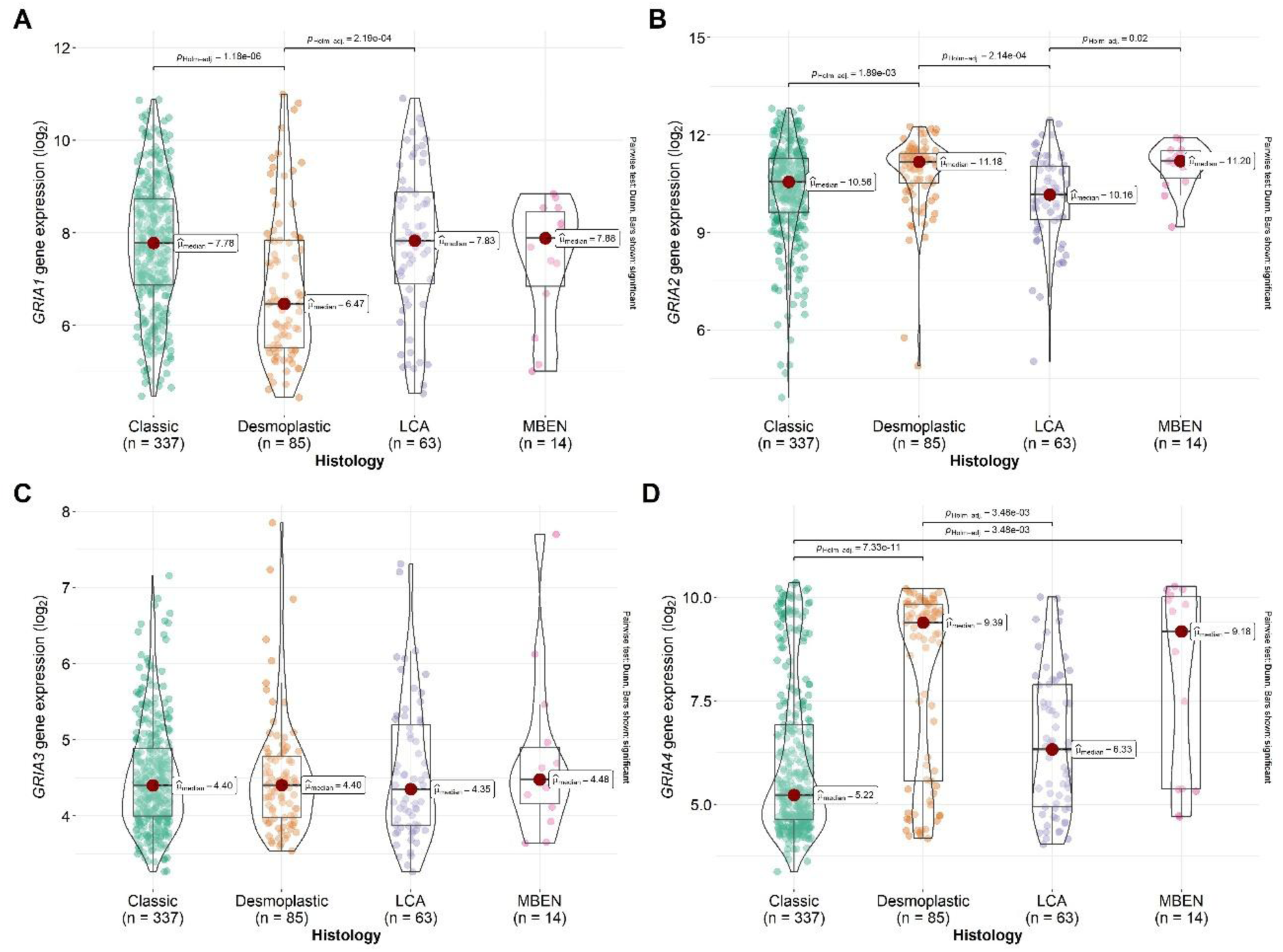
Gene expression levels of **A**, *GRIA1*, **B**, *GRIA2*, **C**, *GRIA3*, and **D**, *GRIA4* in MB tumors classified into histological variants, classic (n = 337), desmoplastic (n = 85), LCA (n = 63), and MBEN (n = 14). Data were obtained from the dataset established by Cavalli et al. (2017). Adjusted *P* values and significant differences are indicated in the panels.

### Expression of *GRIA* Genes in MB Tumors at Single-Cell Resolution

UMAP analysis of *GRIA1-4* expression and distribution among MB subgroups in scRNA-seq of 28 MB samples was consistent with enrichment of *GRIA1* in Group 3 and *GRIA4* in SHH MB (Fig. 4).

**Fig 4.**
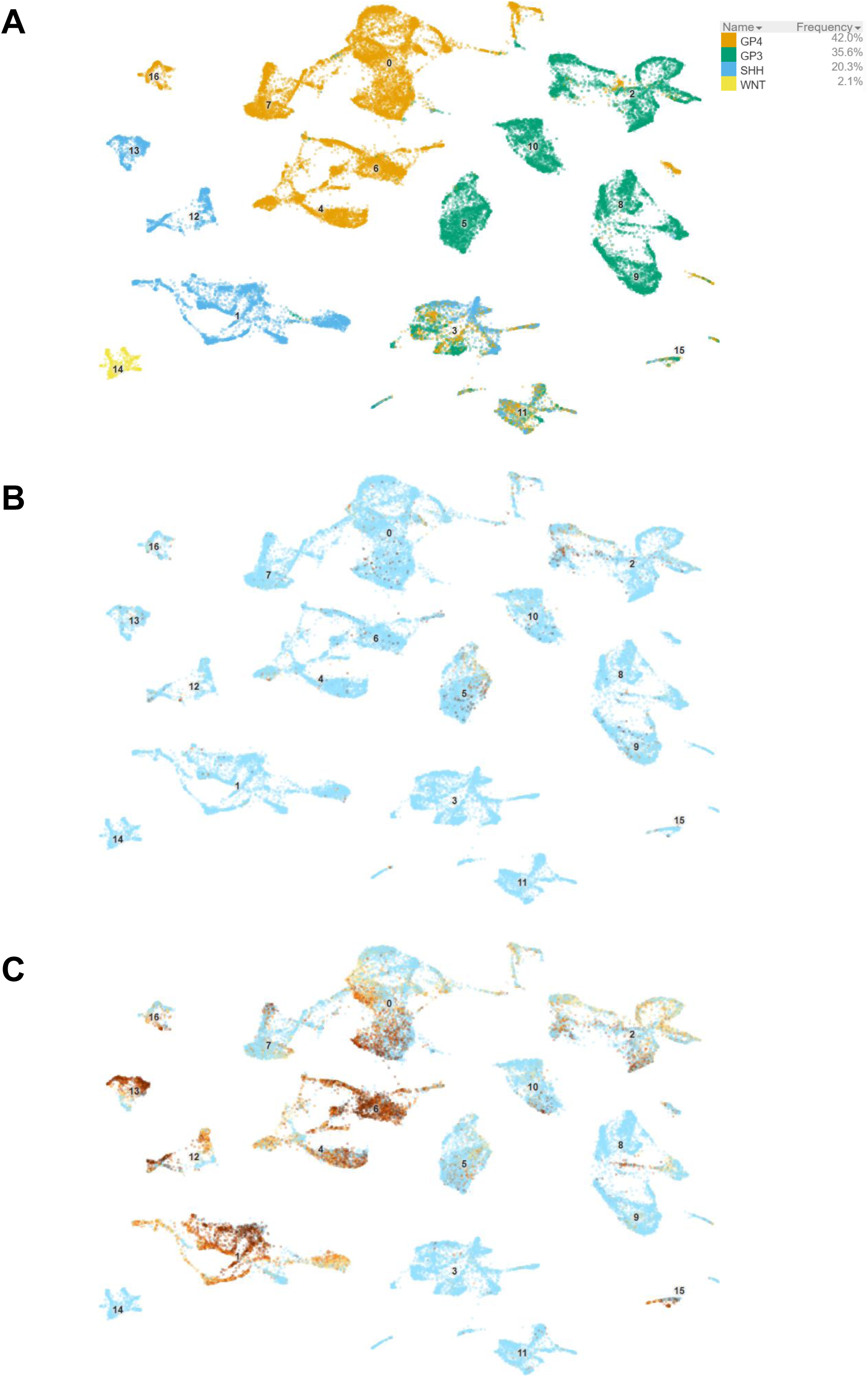
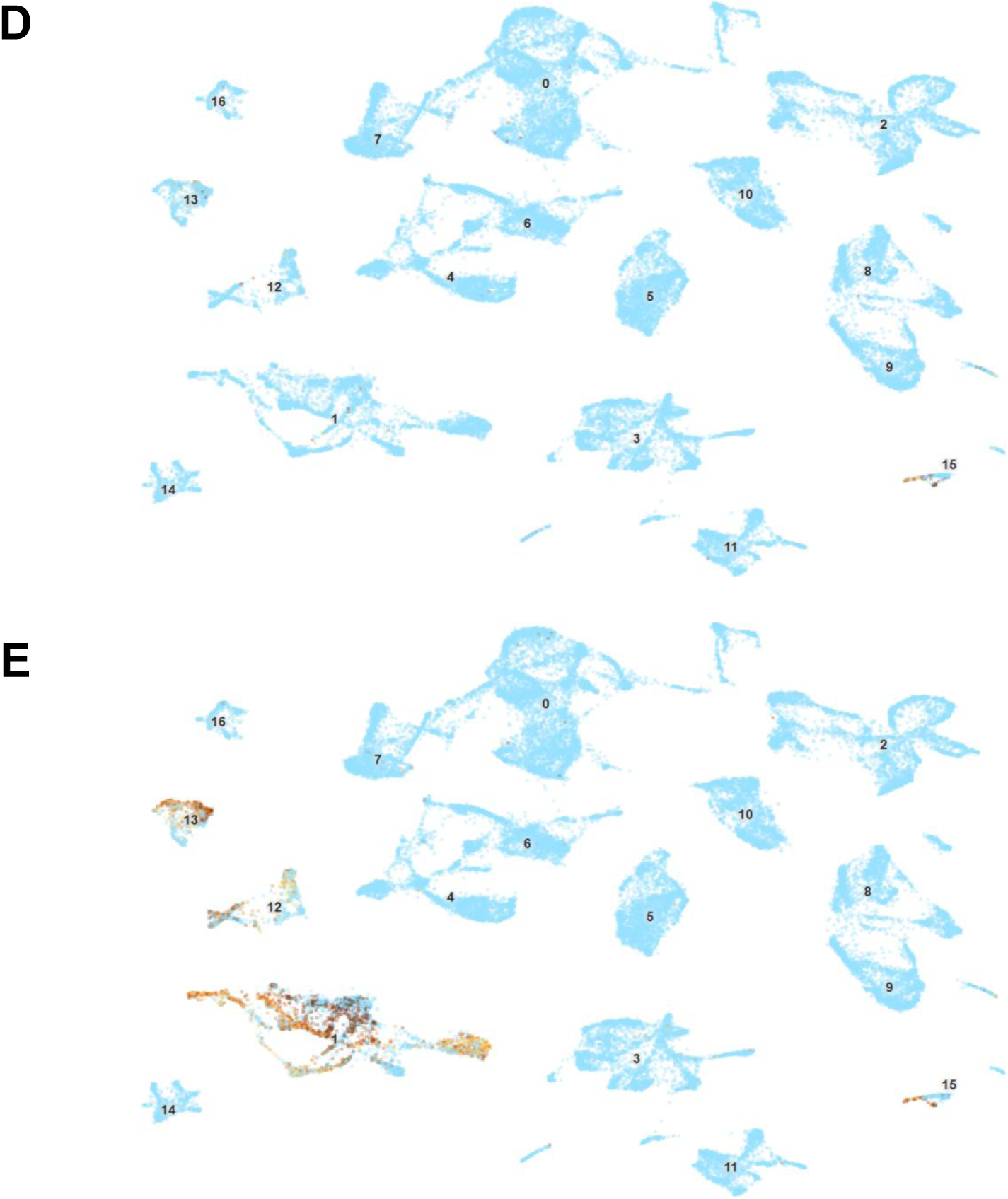
UMAP plots of expression of *GRIA1*-*GRIA4* in the four MB subgroups at single cell resolution. **A,** Cells annotated by molecular subgroup across 28 MB tumor samples. Cells were then annotated by **B,** *GRIA1*, **C,** *GRIA2*, **D,** *GRIA3*, and **E,** *GRIA4*. Data were accessed using single-cell scRNA-seq data available in GSE156053 (Riemondy et al., 2022), as available throught the Pediatric Neuro-oncology Cell Atlas (https://www.pneuroonccellatlas.org/). G3, Group 3 MB; G4, Group 4 MB.

### Expression of *GRIA* Genes Displays Opposing Associations with Patient Survival in Distinct MB Molecular Subgroups

Significant differences in prognosis between tumors with high or low expression of *GRIA1* were observed specifically in Group 3 MB, where higher expression was associated with better survival (Fig. 5). No significant associations with OS were found for *GRIA2* transcription levels (Fig. 6). Patients bearing SHH tumors with high expression of *GRIA3* showed worse survival compared to those with low-expressing tumors (Fig. 6). Finally, for *GRIA4*, high gene expression indicated longer patient OS in WNT and SHH, but worse survival in Group 3 MB (Fig. 8).

**Fig 5.**
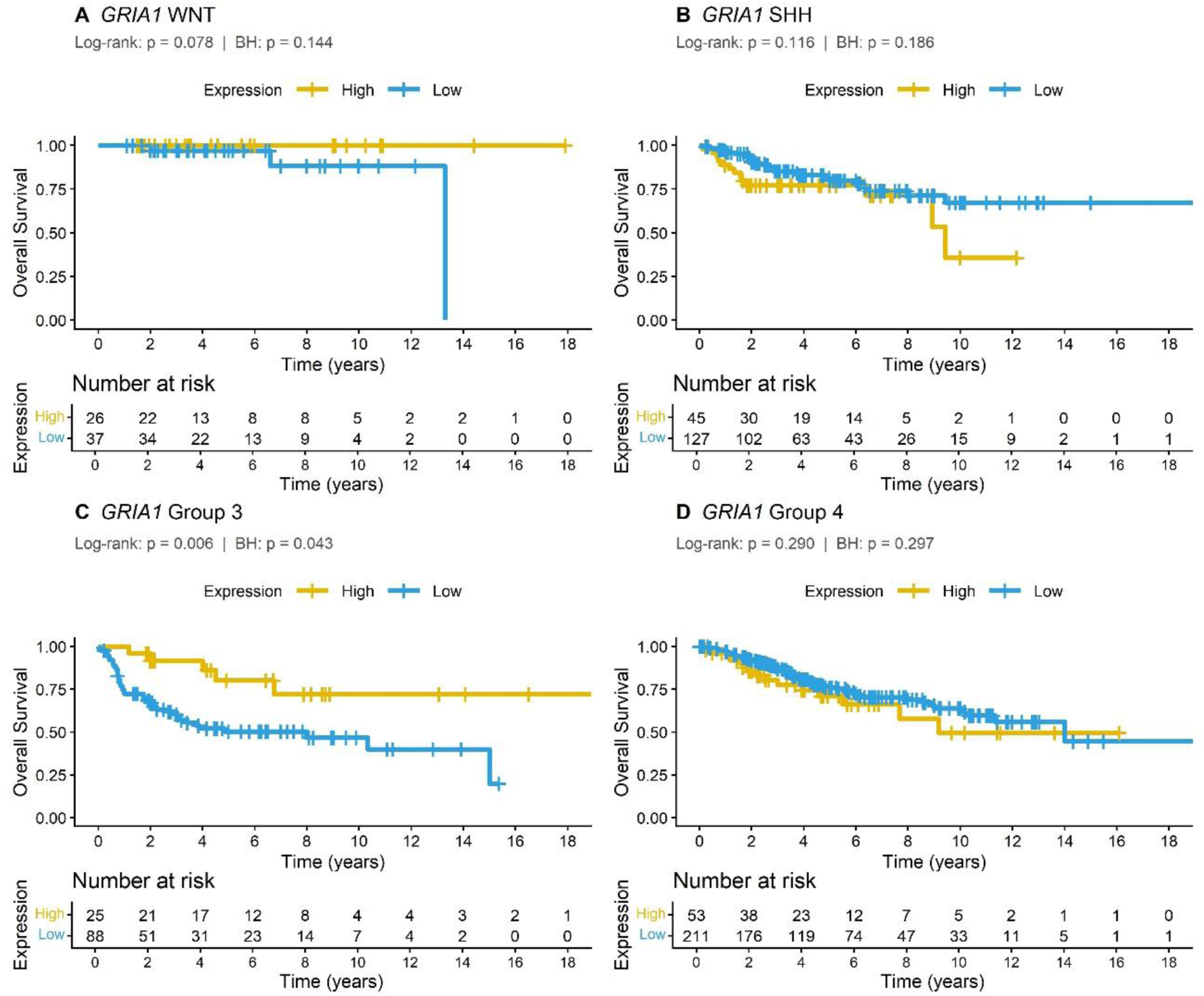
Kaplan-Meier analysis of OS in patients bearing MB tumors with higher or lower expression of the *GRIA1* gene. Tumors were divided into the four molecular subgroups, **A**, WNT (n = 63), **B**, SHH (n = 172), **C**, Group 3 (n = 113), and **D**, Group 4 (n = 264). Data were obtained from the dataset established by Cavalli et al. (2017). Log-rank and adjusted *P* values are indicated in the panels. BH, Benjamini–Hochberg false discovery rate (FDR) correction for multiple hypothesis testing.

**Fig 6.**
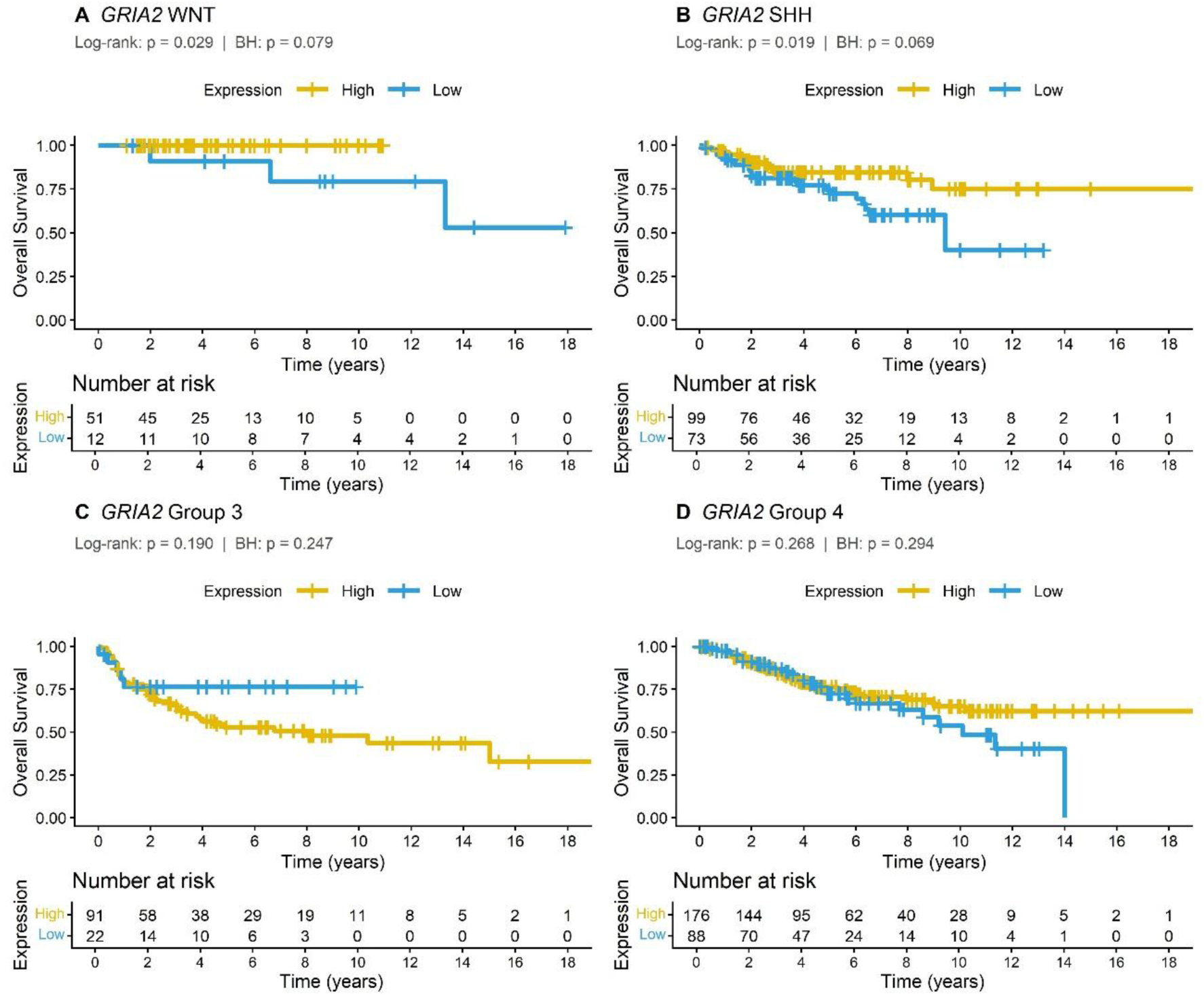
Kaplan-Meier analysis of OS in patients bearing MB tumors with higher or lower expression of the *GRIA2* gene. Tumors were divided into the four molecular subgroups, **A**, WNT (n = 63), **B**, SHH (n = 172), **C**, Group 3 (n = 113), and **D**, Group 4 (n = 264). Data were obtained from the dataset established by Cavalli et al. (2017). Log-rank and adjusted *P* values are indicated in the panels. BH, Benjamini–Hochberg FDR correction for multiple hypothesis testing.

**Fig 7.**
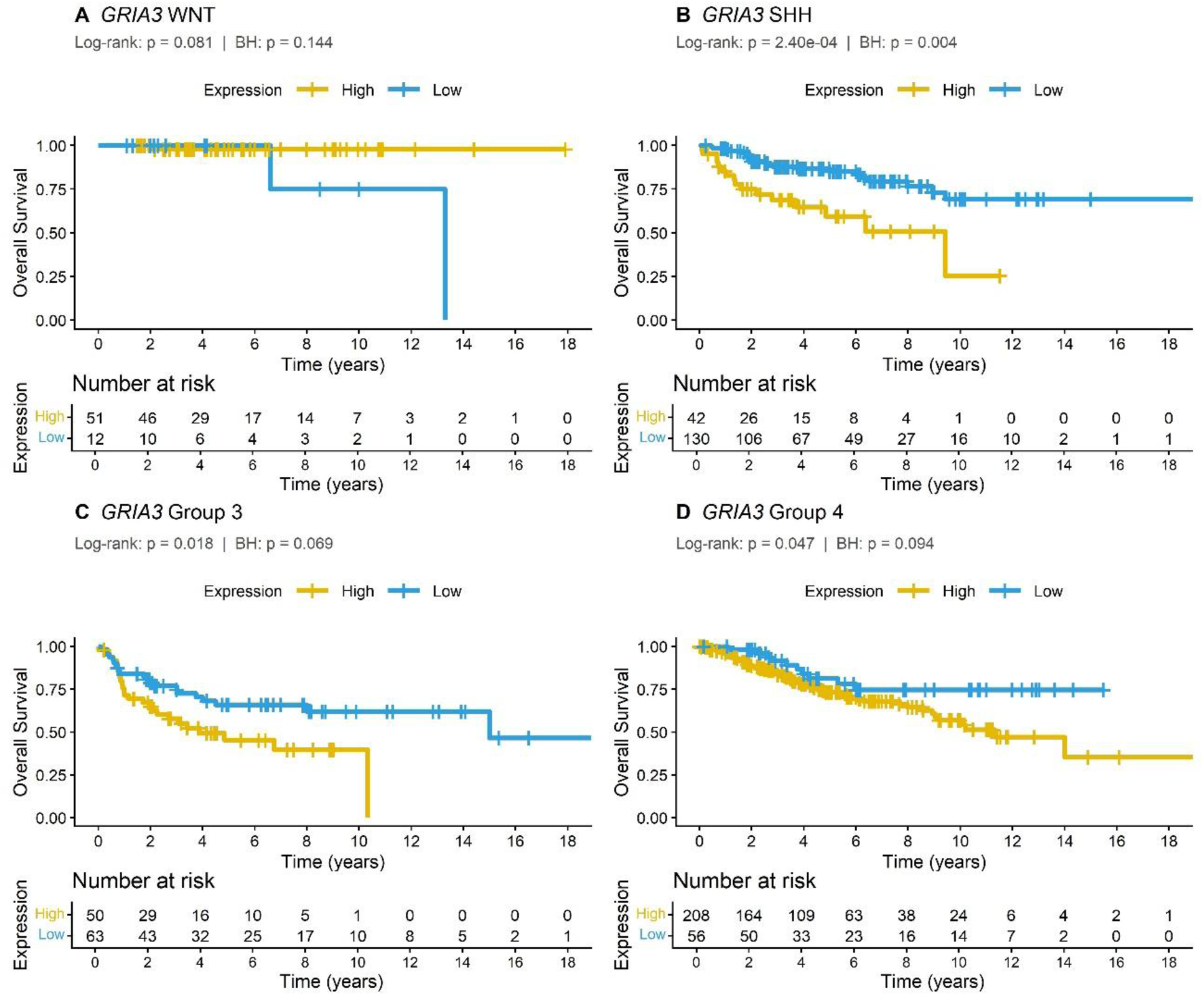
Kaplan-Meier analysis of OS in patients bearing MB tumors with higher or lower expression of the *GRIA3* gene. Tumors were divided into the four molecular subgroups, **A**, WNT (n = 63), **B**, SHH (n = 172), **C**, Group 3 (n = 113), and **D**, Group 4 (n = 264). Data were obtained from the dataset established by Cavalli et al. (2017). Log-rank and adjusted *P* values are indicated in the panels. BH, Benjamini–Hochberg FDR correction for multiple hypothesis testing.

**Fig 8.**
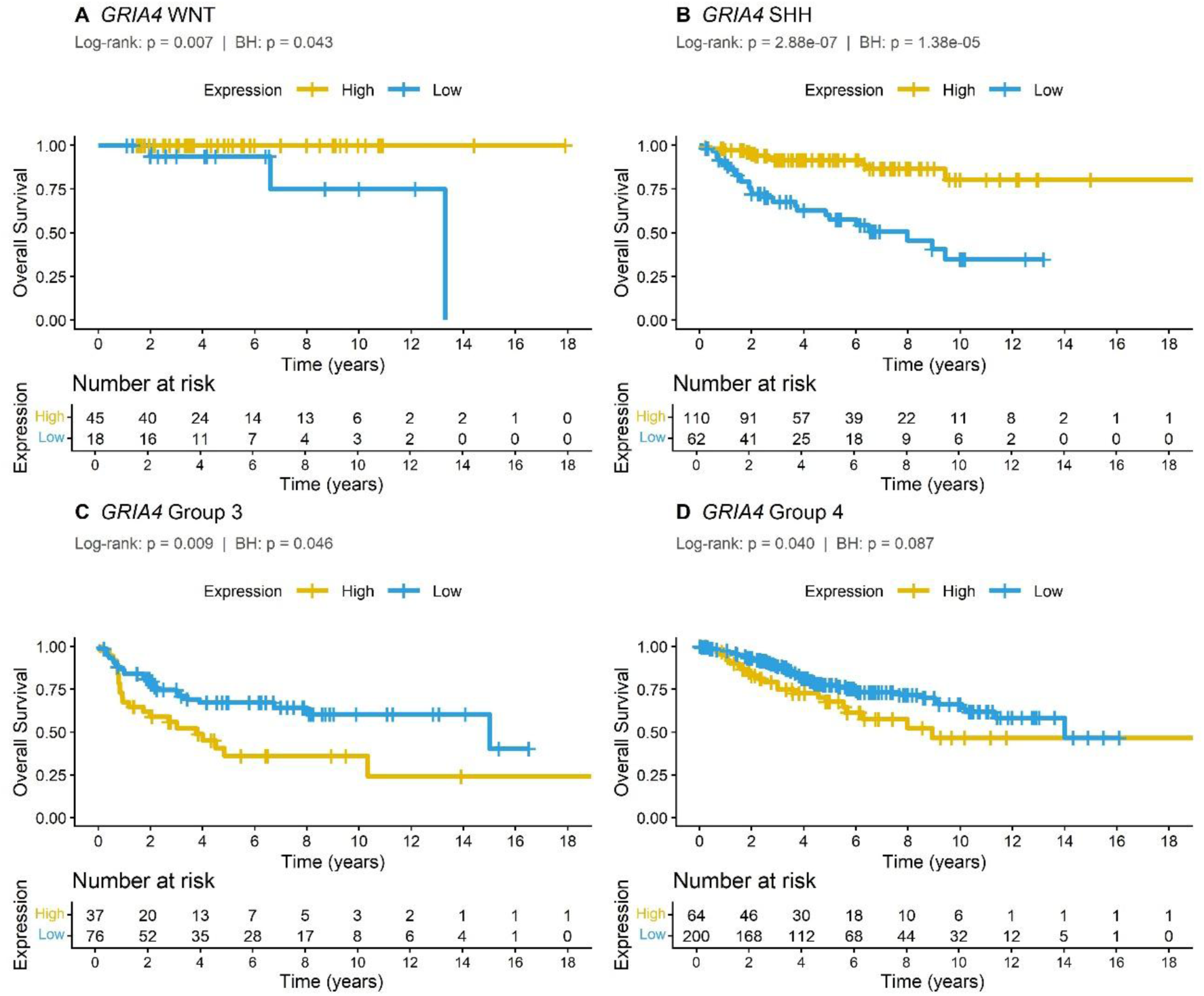
Kaplan-Meier analysis of OS in patients bearing MB tumors with higher or lower expression of the *GRIA4* gene. Tumors were divided into the four molecular subgroups, **A**, WNT (n = 63), **B**, SHH (n = 172), **C**, Group 3 (n = 113), and **D**, Group 4 (n = 264). Data were obtained from the dataset established by Cavalli et al. (2017). Log-rank and adjusted *P* values are indicated in the panels. BH, Benjamini–Hochberg FDR correction for multiple hypothesis testing.

### *GRIA4* Expression is Higher in a SHH MB Cell Line Compared to Group 3/4 and Non-Tumoral Brain Cell Lines

To compare *GRIA*1-4 gene levels in MB cells representative of SHH and Group3/4 MB, we performed RT-qPCR analysis. *GRIA1* and *GRIA2* transcription was significantly higher in D283 cells (representative of Group 3/4 MB) than in DAOY cells (SHH MB), whereas *GRIA4* had much higher levels in DAOY cells (Fig. 9). Consistent with this finding, in cell line data from the Human Protein Atlas, although no formal statistical analysis was applied for comparisons, the value for *GRIA4* expression was visibly higher in DAOY compared to D341 cells (Group 3) and NHAHTDD non-tumoral control brain cells (Supplementary Fig. S1).

**Fig 9.**
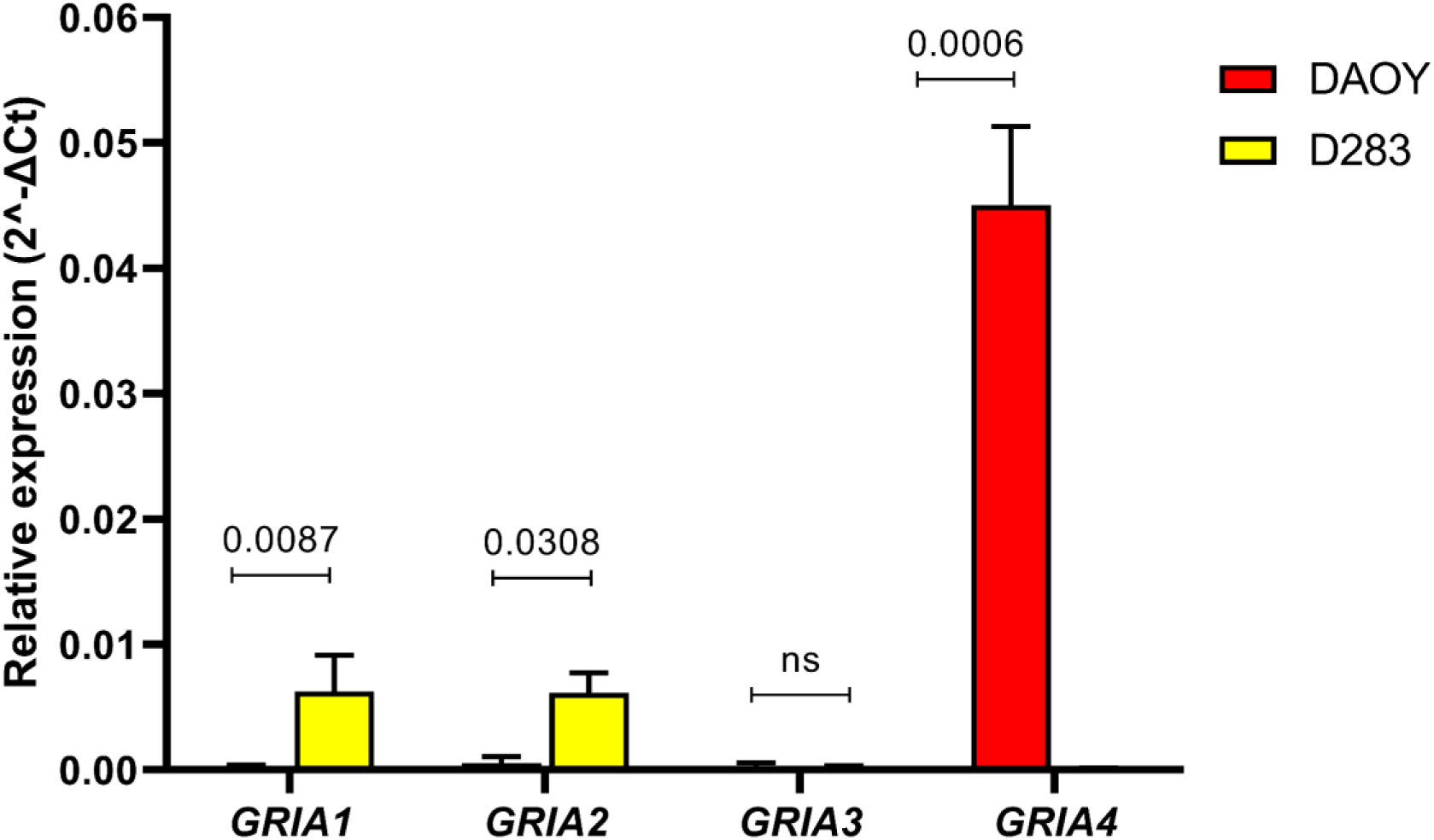
Messenger RNA expression of *GRIA1-4* in DAOY and D283 MB cell lines. Analysis was performed using RT-qPCR as described above; *P* values are shown in the panels.

### Associations of Genes Encoding NMDA and Kainate Receptor Subunits with OS in MB Patients

We extended our analyses of potential prognostic value to a selected set of genes encoding core functional subunits that define canonical NMDA and kainate receptor signaling, namely *GRIN1* together with *GRIN2A* and *GRIN2B*, which constitute the obligatory and principal regulatory NMDA receptor subunits, and *GRIK1–4*, which encode the main pore-forming kainate receptor subunits capable of mediating ionotropic signaling (Paoletti P. et al., 2013). High expression of *GRIN2A* was associated with longer, and of *GRIN2B* with shorter, OS in SHH MB (Supplementary Fig. S2). For kainate receptor subunit genes, high expression of *GRIK3* in WNT or of *GRIK4* in Group 3 tumors, indicated better survival (Supplementary Fig. S2).

## Discussion

AMPARs on glioma cell membranes have been proposed to mediate glutamatergic transmission at synapses formed between the tumors and surrounding neurons, which are hijacked to become part of the microenvironment system supporting cancer progression (Taylor et al., 2023; Venkataramani et al., 2019; Venkatesh et al., 2019). In contrast, the structural and functional associations between brain tumor cells and non-tumoral brain tissue remain much less understood in MB. The guiding hypothesis of the present study is that expression of AMPAR subunits in MB tumors, and their association with patient survival, may indicate that MB, similar to gliomas, recruits synaptic transmission and neural plasticity mechanisms.

Previous reports have shown that the expression of candidate prognostic genes can have very different patterns of association with patient survival across the distinct molecular subgroups of MB (Coelho et al., 2025; Monteiro et al., 2024; Thomaz et al., 2020; Park et al., 2019). The present findings are consistent with that view, given that expression of *GRIA* genes showed bidirectional and subgroup-dependent associations with prognosis, with individual subunits linked to either improved or worse survival depending on molecular context. *GRIA3* and *GRIA4* showed particularly strong associations with survival in SHH MB, with each gene demonstrating effects in opposite directions. Analyses of selected NMDA and kainate receptor subunit genes showed opposite patterns of association of *GRIN2A* and *GRIN2B* in SHH tumors, and better prognosis with higher expression of *GRIK3* in WNT and *GRIK4* in Group 3 MB. These additional receptor subunits were analyzed to determine whether prognostic associations observed for AMPAR genes extend to other ionotropic glutamate receptor families involved in excitatory synaptic transmission. The results also underscore the crucial importance of stratifying MB tumors by molecular subgroup when evaluating candidate prognostic biomarkers.

Given that AMPAR activity is proposed to contribute to brain cancer progression, the findings that expression of three *GRIA* genes is lower in MB tumors than normal cerebellar tissue, and that high expression of *GRIA* genes in several MB subgroups indicates a more favorable prognosis, are unexpected. Nevertheless, studies in malignant gliomas suggest that AMPAR function is more nuanced than a purely pro-tumorigenic role. Sustained receptor-mediated calcium influx may lead to intracellular calcium overload, triggering excitotoxic mechanisms and subsequent tumor cell injury (Radin et al., 2025; Liu et al., 2015). In line with this, pharmacological activation of AMPARs has been reported to decrease glioblastoma cell viability and induce apoptotic responses in other cancer types (Radin et al., 2018). Together, these observations raise the possibility, albeit speculative at present, that under certain conditions AMPAR activity may exert antitumoral, rather than oncogenic, effects. The subgroup-dependent and often opposing associations between *GRIA* expression and survival in MB suggest that AMPAR subunits may reflect underlying lineage-specific differentiation states or neuron-like transcriptional programs rather than exerting a uniform pro- or anti-tumorigenic effect. It is also possible that increased AMPAR content may indicate a more mature neuron-like phenotype. Promoting neuronal differentiation in glioma or MB cells can reduce cell proliferation and expression of tumor-associated proteins, while upregulating tumor suppression genes (Yi et al., 2024).

Cell lines are widely used as experimental models for exploring aspects of MB biology and raising novel hypotheses about candidate therapeutic targets. DAOY cells represent SHH, *TP53*-mutated MB, whereas D283 and D341 cells are classified as Group 3 MB or Gorup3/4 (Casciati et al., 2020; Friedman et al., 1988; Ivanov et al., 2016; Jacobsen et al., 1985). Our analysis of both RT-qPCR results and data available in The Human Protein Atlas suggested a pattern that is overall consistent with a markedly higher expression of *GRIA4* in SHH tumors compared to other MB subgroups, as well as with the differences we observed between SHH and Group 3 MB at tumor sample and single-cell levels. Thus, these findings support the validity of these cell lines as surrogate models to investigate the biology of different MB subgroups.

Our study has clear limitations. Links between gene expression and patient outcome do not necessarily point to a direct mechanistic or causal role. Also, it is well known that higher mRNA expression does not always translate into higher protein levels. While some studies in different model systems have reported a reasonable correspondence between mRNA and protein content, others have found only weak correlations (Greenbaum et al., 2002; Pevsner, 2015; Waters et al., 2006). Further work will therefore be needed to directly assess the expression of AMPAR subunit proteins and to clarify their possible roles in MB.

## Conclusion

We describe the mRNA expression of *GRIA* genes in MB tumors and MB cell lines stratified according to different molecular subgroups or histological variants, and the specific patterns of association between expression levels of each of the distinct *GRIA* genes with patient outcome. Future studies should evaluate AMPAR protein expression and functional activity in MB models to determine whether AMPAR signaling contributes directly to tumor biology or reflects lineage-specific differentiation states.

## Supporting information

Supplementary Information-Saciloto et al

## Author Contributions

BS, RR, and GRI conceived the study. BS, MD, JCM, ML, JV, IBSR, and AT conducted analyses and experiments. MS, JCM, JV, IBSR, and AT plotted the data. BS, MD, JCM, ML, JV, JMRF, CBF, AT, RR, and GRI interpreted the results. BS, MD, and RR wrote the first draft of the manuscript. BS, OM, MACF, CBF, RR, and GIR provided funding and resources. MACF, AT, RR, and GIR supervised the study. BS, MD, IBSR, JMRF, OM, MACF, AT, RR, and GIR revised the final version of the manuscript.

## Data Availability

The tumor datasets used in this study are available in the Gene Expression Omnibus (GEO) repository, accession number GSE202043, https://www.ncbi.nlm.nih.gov/geo/query/acc.cgi?acc=GSE202043; accession number GSE85217, https://www.ncbi.nlm.nih.gov/geo/query/acc.cgi?acc=GSE85217, and GSE156053 (https://www.ncbi.nlm.nih.gov/geo/query/acc.cgi?acc=GSE156053. The MB scRNA-seq single cell dataset is available at the Pediatric Neuro-oncology Cell Atlas (https://www.pneuroonccellatlas.org/). Data from cell lines were obtained from The Human Protein Atlas (https://www.proteinatlas.org/, accessed on January 27^th^ 2026) and are available at https://www.proteinatlas.org/ENSG00000155511-GRIA1/cell+line#brain_cancer; https://www.proteinatlas.org/ENSG00000120251-GRIA2/cell+line#brain_cancer; https://www.proteinatlas.org/ENSG00000125675-GRIA3/cell+line#brain_cancer; https://www.proteinatlas.org/ENSG00000152578-GRIA4/cell+line#brain_cancer; https://www.proteinatlas.org/ENSG00000155511-GRIA1/cell+line#non-cancerous; https://www.proteinatlas.org/ENSG00000120251-GRIA2/cell+line#non-cancerous; https://www.proteinatlas.org/ENSG00000125675-GRIA3/cell+line#non-cancerous; and https://www.proteinatlas.org/ENSG00000152578-GRIA4/cell+line#non-cancerous.

## Declarations

### Ethical Approval and Consent to Participate

Not applicable.

### Consent for Publication

Not applicable.

## Abbreviations

AMPA: α-amino-3-hydroxy-5-methyl-4-isoxazolepropionic acid
AMPAR: AMPA receptor
BH: Benjamini–Hochberg
FDR: False discovery rate
G3: Group 3 medulloblastoma
G4: Group 4 medulloblastoma
GBM: Glioblastoma
GCP: Granule cell precursor
GEO: Gene Expression Omnibus
GNP: Granule neuron precursor
LCA: Large cell/anaplastic
MB: Medulloblastoma
MBEN: Medulloblastoma with extensive nodularity
mRNA: Messenger ribonucleic acid
NS: Non-significant
OS: Overall survival
qRT-PCR: Quantitative reverse transcription polymerase chain reaction
RL: Rhombic lip
scRNA-seq: Single-cell RNA sequencing
SHH: Sonic hedgehog
Sig.: Significant difference
UMAP: Uniform Manifold Approximation and Projection
WNT: Wingless

