## Supplementary Information-Saciloto et al for "Subgroup-Specific Associations of *GRIA* Genes Encoding AMPA Glutamate Receptor Subunits with Patient Survival in Medulloblastoma"

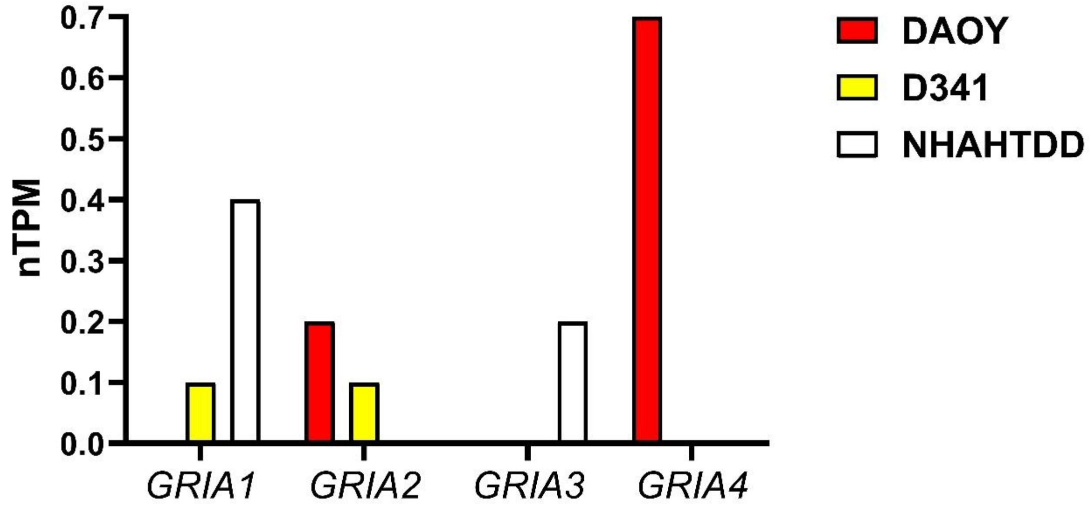


**Supplementary Fig S1** Messenger RNA (expression of *GRIA* genes in DAOY MB, D341 MB, and NHAHTDD non-tumoral brain cells. Data normalized as transcript per million (nTMP) were obtained from The Human Protein Atlas (https://www.proteinatlas.org/; accessed on January 27th 2026).

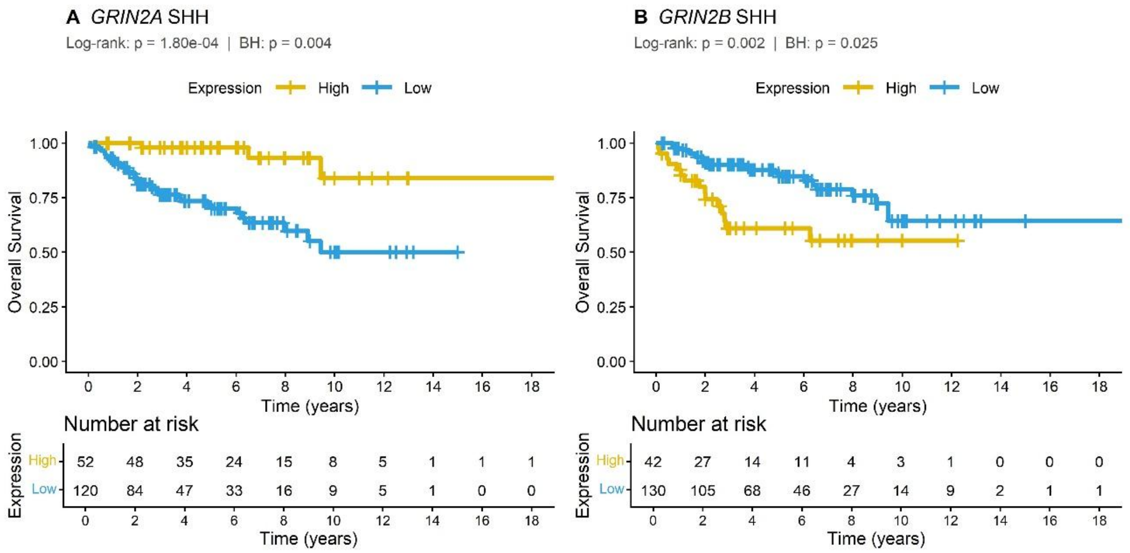


**Supplementary Fig S2** Kaplan-Meier analysis of OS in patients bearing SHH MB tumors (n = 172) with higher or lower expression of the **A,** *GRIN2A*, and **B,** *GRIN2B* genes. Data were obtained from the dataset established by Cavalli et al. (2017). Log-rank and adjusted *P* values are indicated in the panels. BH, Benjamini–Hochberg FDR correction for multiple hypothesis testing.

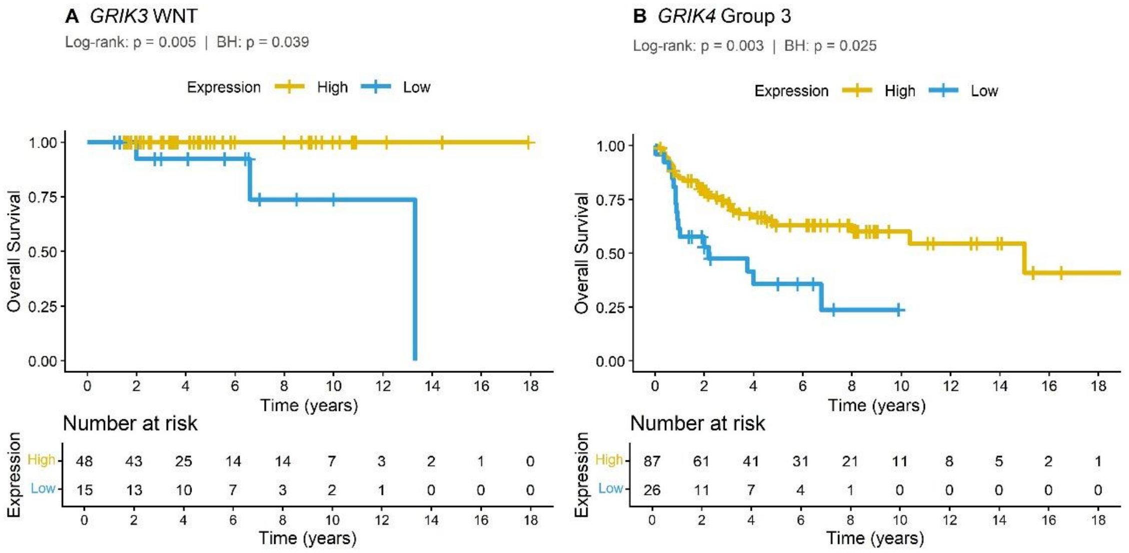


**Supplementary Fig S3** Kaplan-Meier analysis of OS in patients bearing **A,** WNT MB tumors (n = 63) with higher or lower expression of the *GRIK3* gene, and **B,** Group 3 MB tumors (n = 113) with higher or lower expression of the *GRIK4* gene. Data were obtained from the dataset established by Cavalli et al. (2017). Log-rank and adjusted *P* values are indicated in the panels. BH, Benjamini–Hochberg FDR correction for multiple hypothesis testing.
